# Highly Thermotolerant SARS-CoV-2 Vaccine Elicits Neutralising Antibodies Against Delta and Omicron in Mice

**DOI:** 10.1101/2022.03.03.481940

**Authors:** Petrus Jansen van Vuren, Alexander J. McAuley, Michael J. Kuiper, Nagendrakumar B. Singanallur, Matthew P. Bruce, Shane Riddell, Sarah Goldie, Shruthi Mangalaganesh, Simran Chahal, Trevor W. Drew, Kim R. Blasdell, Mary Tachedjian, Leon Caly, Julian D. Druce, Shahbaz Ahmed, Mohammad Suhail Khan, Sameer Kumar Malladi, Randhir Singh, Suman Pandey, Raghavan Varadarajan, Seshadri S. Vasan

## Abstract

As existing vaccines fail to completely prevent COVID-19 infections or community transmission, there is an unmet need for vaccines that can better combat SARS-CoV-2 variants of concern (VOC). We have previously developed highly thermo-tolerant monomeric and trimeric receptor binding domain derivatives that can withstand 100°C for 90 minutes and 37°C for four weeks, and help eliminate cold chain requirements. We show that mice immunised with these vaccine formulations elicit high titres of antibodies that neutralise SARS-CoV-2 variants VIC31 (with Spike: D614G mutation), Delta and Omicron (BA.1.1) VOC. Compared to VIC31, there was an average 14.4-fold reduction in neutralisation against BA.1.1 for the three monomeric antigen-adjuvant combinations, and 16.5-fold reduction for the three trimeric antigen-adjuvant combinations; the corresponding values against Delta were 2.5 and 3.0. Our findings suggest that monomeric formulations are suitable for the upcoming Phase I human clinical trials, and that there is potential for increasing efficacy with vaccine matching to improve responses against emerging variants. These findings are consistent with *in silico* modelling and AlphaFold predictions which show that while oligomeric presentation can be generally beneficial, it can make important epitopes inaccessible, and also carries the risk of eliciting unwanted antibodies against the oligomerisation domain.

## 1. Introduction

The novel coronavirus disease 19 (COVID-19) was declared by the World Health Organization (WHO) as a ‘Public Health Emergency of International Concern’ on 30 January 2020 and as a pandemic on 11 March 2020. The WHO originally predicted that COVID-19 would take 4-5 years to control [1], and as we enter the third year of the on-going pandemic, this target seems tantalisingly close. With already 428 million cases and 5.91 million deaths as of 24 February 2022 [2], vaccination remains the key defence against the severe acute respiratory syndrome coronavirus 2 (SARS-CoV-2).

In the context of vaccination, the WHO has listed four factors - poor coverage (especially in vulnerable populations), inequitable access, sub-optimal duration of protection post-vaccination (against infection, severe disease and death), and the emergence of variants of concern (VOC) – as the key ones driving the impact of SARS-CoV-2 [3]. The first two factors, poor coverage and inequitable access, are interlinked and impacted by cold chain storage requirements, while the next two factors, sub-optimal protection and VOC, are also interlinked.

Whilst 10.6 billion doses of the COVID-19 vaccines have been administered globally to date, indicating that 4.4 billion of the world’s population is fully vaccinated [2]; the reality is that half the world’s population, predominantly in low income countries (LICs) and lower middle-income countries (LMICs), is still receiving its first dose of a COVID-19 vaccine, whilst administration of the third dose is well underway in most upper middle income countries (UMICs) and high income countries (HICs), with Israel having already giving the fourth dose for its nationals [4]. LICs have administered 13-fold fewer vaccines compared to both UMICs and HICs, while LMICs (including India which is responsible for 62% of the global vaccine production) are below the world-average (**Figure 1A**) [5].

**Figure 1.**
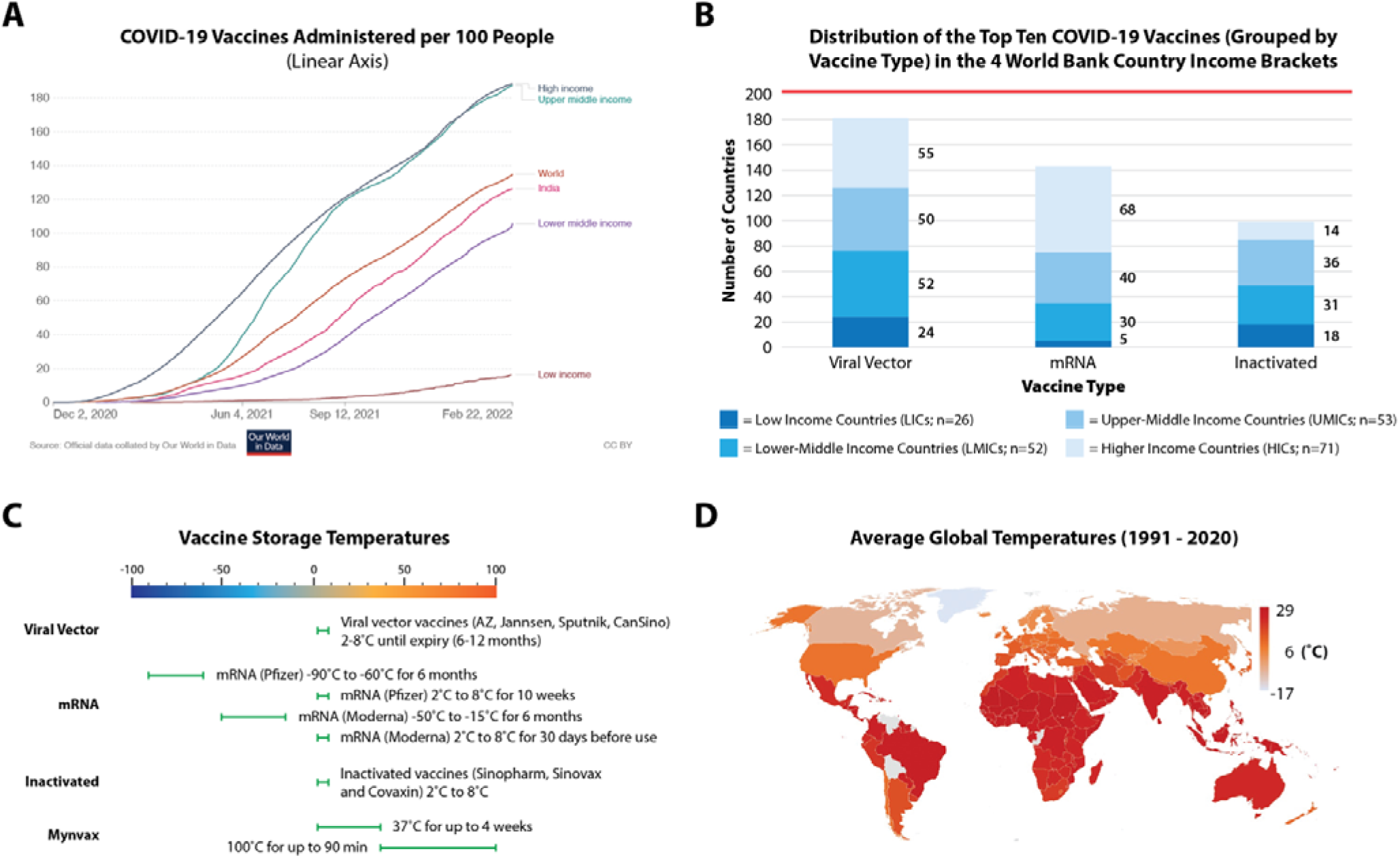
COVID-19 vaccine storage temperature and cold chain affects equitable access. Total doses (including boosters) administered per 100 people is shown in **(A)**, while **(B)** shows the top 10 vaccines (grouped into 3 categories) inequitably distributed across the 4 World Bank country income brackets. **(C)** Storage temperatures for these top 10 vaccines (grouped under 3 headings) is compared to Mynvax formulation. **(D)** Average storage temperatures show a high level of correlation with vaccine inequity experienced by LICs and LMICs. In **B**, the red line denotes the total number of countries used in our analysis (n = 202) based on availability of information from Our World in Data. The data in **A** was accurate as of 22 February 2022, and **B** accurate as of 6 February 2022.

Grouping the world’s top 10 most-administered vaccines based on their type – a) Viral Vector vaccines [AstraZeneca/AZ (Vaxzevria/AZD1222), Janssen, SputnikV/Sputnik Light (Gam-COVID-Vac), Convidecia/CanSino (AD5-nCOV)], b) mRNA vaccines [Pfizer (Comirnaty/BNT162b2), Moderna (Spikevax/mRNA-1273)], and c) Inactivated vaccines [Sinopharm (BBIBP-CorV/ NVSI-06-07), Covaxin (BBV152), Sinovac (CoronaVac)] – we see that the viral vectored and inactivated vaccines (which only require storage at 2-8°C) are more evenly distributed across the different country income brackets (**Figures 1B and 1C**). In contrast, Pfizer requires ultra-cold-chain freezer capacity at −70°C, and Moderna requires traditional freezer capacity at −20°C (**Figure 1C**); these have largely benefited HICs and UMICs, which also have greater manufacturing expertise for mRNA vaccines, highlighting that vaccine inequity and cold storage requirements are highly correlated.

As no country is protected from this virus until all countries are, it is of paramount importance to address this inequity, keep up with the emergence of new variants, and improve vaccination rates in LICs and LMICs. These countries are ill-equipped for the financial and logistical burdens of cold chain transport and storage requirements, which are necessary for all approved COVID-19 vaccines to varying degrees (**Figure 1C**). This is one of the key contributing factors disproportionately impacting LICs and LMICs where the average temperatures are higher (**Figure 1D**) [6,7].

At 14.2 doses per 100 people, most people in LICs haven’t received their first dose; we have ample evidence that a single dose does not adequately prevent infections both at an individual level and community transmission. For instance, with the Pfizer vaccine, protection against symptomatic COVID-19 was only 52% twelve days after the first dose [8]. This problem has been exacerbated by the emergence of variants - five of which have been declared to be of concern and newer ones could arise in poorly vaccinated regions. The duration of protection is even lower with the recent Delta and Omicron; our recent study with AstraZeneca, Moderna and Pfizer shows that at least a third dose is required to generate sufficient neutralising antibody titres against these two VOCs [9]. Therefore, we urgently need next-generation vaccines (as described in this paper) which are not only highly effective against all existing and emerging variants of SARS-CoV-2 but are also thermostable and do not have cold chain transport and storage requirements.

## 2. Materials and Methods

In this paper, we analyse the serological responses of the VOCs in mice for six vaccine formulations developed by Mynvax Private Limited, one of which is to be selected for upcoming Phase I human clinical trials. The underlying antigens are highly thermo-tolerant monomers (Vaccines 1-3, c.f. §2.1) or trimers (Vaccines 4-6, c.f. §2.2) as shown in **Figure 2**. Mynvax has previously shown that its receptor binding domain derivatives can withstand temperatures up to 100°C for 90 minutes and 37°C for four weeks [10–12], therefore this paper will not go into proving thermostability. Details of mice immunisation are presented in §2.3, while §2.4-2.5 describe the materials and methods for live virus neutralisation assays. §2.6 and 2.7 respectively describe the *in silico* and statistical methods used to interpret results.

**Figure 2.**
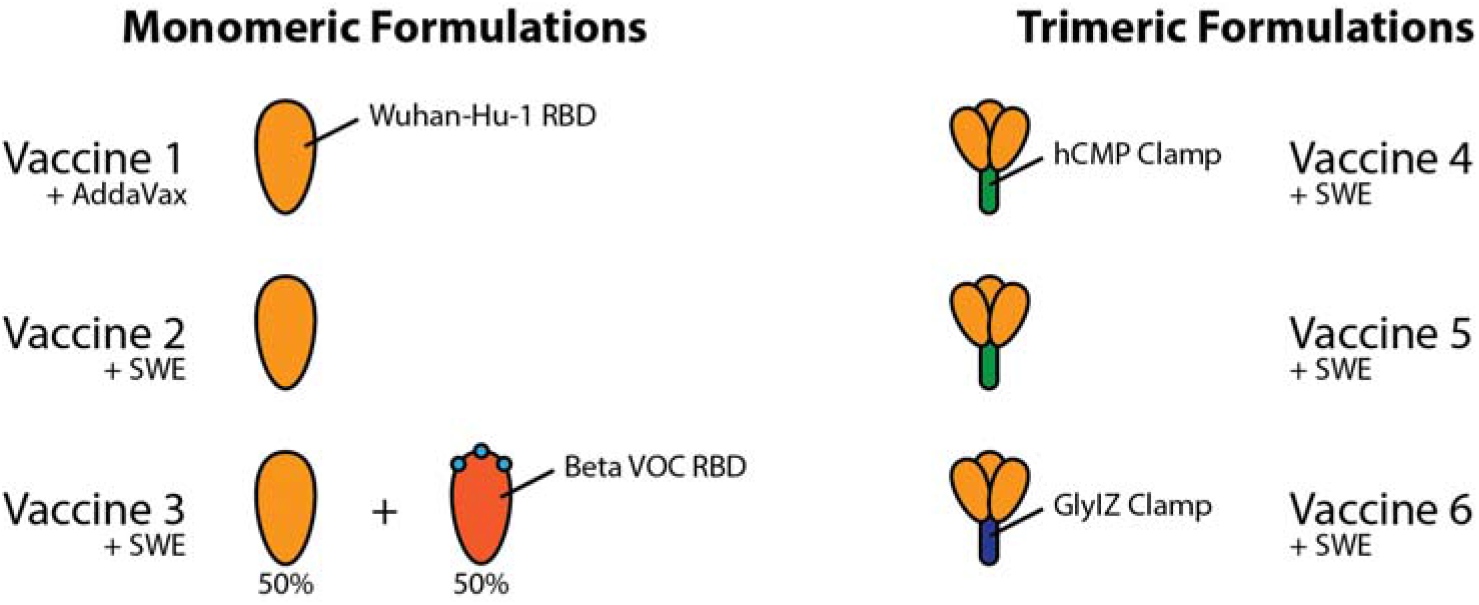
Schematic diagram of vaccine formulations 1-6 evaluated by this study (c.f. **Table 1**). **Supplementary Text A** contains further amino acid information.

### 2.1. Thermostable Monomer-Adjuvant Formulations

The RBD of SARS-CoV-2 and its mutants were cloned in the pcDNA3.4 vector for expression in Expi293F cells as described earlier [10,11]. After transfection and protein purification, the mRBD1-3.2 RBD protein was obtained at a yield of ~180 mg/l. These mammalian cell-expressed wild-type (WT) and mRBD1-3.2 RBD at 0.2 mg/ml of PBS or adjuvants (SWE) were stored at 4°C and 45°C for up to 28 days. Aliquots were taken at regular intervals and diluted in PBS to a final concentration of 100 nM, and the amount of folded protein remaining was estimated using surface plasmon resonance (SPR). In another set of experiments, following dialysis against water and lyophilisation, RBD was subjected to thermal incubation at 37°C for up to 30 days in individual aliquots. At each time point, aliquots were returned to 4°C. Prior to SPR and differential scanning fluorimetry (DSF), samples were resolubilised in PBS at concentrations of 100 nM and 0.2 mg/ml, respectively. SPR binding to immobilised angiotensin-converting enzyme 2 (ACE-2) hFc and DSF were performed as described previously.

The following ‘Mynvax’ protein sub-unit monomer vaccine formulations 1-3 were prepared, as follows:

Vaccine 1: mRBD1-3.2 is a stabilised multi-mutant of RBD (residues 331-532) of SARS-CoV-2 Spike (S) protein. It contains A348P, Y365W and P527L stabilising mutations [10,11]. The protein was expressed in mammalian Expi293F cells (Thermo Fisher Scientific, Cat no. A14527) and purified using Ni-NTA (GE Healthcare, Cat no. 17531802) affinity purification. This antigen has been shown to be stable at 37°C for up to a month without any reduction in the amount of folded fraction. The protein antigen was adjuvanted with AddaVax (1:1 v/v; vac-adx-10, InvivoGen, USA) for immunisation.

Vaccine 2: The stabilised antigen mRBD1-3.2 was adjuvanted with a squalene-in-water emulsion adjuvant (SWE), a GMP grade adjuvant equivalent to MF59 (Sepivac SWE, 1:1 v/v; Cat. No. 80748, SEPPIC SA, France) for immunisation.

Vaccine 3: A formulation of equal amounts (20 μg each) of mRBD1-3.2 and mRBD1-3.2-beta was adjuvanted with SWE (1:1 v/v) for immunisation. mRBD1-3.2-beta was generated in the background of stabilised mRBD1-3.2 (A348P, Y365W and P527L) and it has three important mutations (K417N, E484K and N501Y) present in the RBD of the Beta (B.1.351) VOC.

### 2.2. Trimeric Human Cartilage Matrix Protein (hCMP) Formulations

Malladi et al. [10] previously designed a monomeric glycan engineered derivative of the receptor binding domain termed mRBD (residues 332-532 possessing an additional glycosylation site at N532) that induced neutralising antibodies in guinea pig immunisations. Oligomerisation of native antigens was expected to induce higher titres of binding and neutralising antibodies, therefore the mRBD was fused to the disulphide linked trimerisation domain derived from hCMP (residues 298–340). We hypothesised that RBD fused to the hCMP trimerisation domain (residues 298–340) would elicit higher neutralising antibody titres relative to the corresponding monomer. The ‘Mynvax’ protein sub-unit trimer vaccine formulations 4-6 used in this work were as follows:

Vaccine 4: The WT RBD (residues 332-532) was fused to the C-terminus of the disulphide linked trimerisation domain derived from hCMP and the resultant construct was called hCMP-mRBD [12]. The protein was expressed in mammalian Expi293F cells (Thermo Fisher Scientific, Cat no. A14527) and purified using Ni-NTA (GE Healthcare, Cat no. 17531802) affinity purification. This antigen was adjuvanted with SWE (1:1 v/v) for immunisation. This antigen was shown to be highly thermotolerant and remained folded for up to a month when lyophilised powder was incubated at 37°C, the protein was also tolerant to transient thermal stress to 100°C for up to 90 minutes.

Vaccine 5: The above mentioned hCMP-mRBD antigen when expressed from Stable CHO (Thermo Fisher Scientific, Cat no. R75807) cell lines was named as hCMP-mRBD-CHO. This antigen was adjuvanted with SWE (1:1 v/v) for immunisation.

Vaccine 6: The glycosylated synthetic trimerisation domain IZN4 [13] was fused at the C-terminus of WT RBD (residues 332-532) [12]. The presence of glycosylation in the trimerisation domain reduces the immune response against the scaffold. This antigen was adjuvanted with SWE (1:1 v/v) for immunisation.

### 2.3. Mouse immunisation

The purified protein antigens (20 μg per animal) were formulated with the ‘MF59’ equivalent squalene-based oil-in-water emulsion AddaVax or SWE adjuvants. The formulations were administered to 6–8-week-old female BALB/c mice (n=5) via intra-muscular injection, on Day 0 (prime), Day 21 (first boost), and Day 42 (second boost). The protein antigen (20 μg in 50 μl) was diluted with the adjuvant (50 μl) before immunisation. Pre-bleed serum was collected two days before immunisation, and two weeks post prime and boosts, serum was collected to estimate the titres of elicited IgG antibodies. The study was conducted at Central Animal Facility, Indian Institute of Science, Bangalore, India. All animal studies were approved by the Institutional Animal Ethics Committee (IAEC no. CAF/ETHICS/799/2020). The mouse samples were imported into ACDP, Geelong, Australia and were gamma irradiated on entry per import permit conditions.

### 2.4. SARS-CoV-2 isolation and stocks

Three SARS-CoV-2 isolates viz., VIC31-D614G (hCoV-19/Australia/VIC31/2020, containing the D614G mutation) and the two variants of concern (VOC) Delta (hCoV-19/Australia/VIC18440/2021), and Omicron BA.1.1 (hCoV-19/Australia/VIC28585/2021), were kindly provided by Drs Caly and Druce at the Victorian Infectious Diseases Reference Laboratory (VIDRL; Melbourne, Australia). Virus stocks were propagated and titrated in Vero E6 cells (American Type Culture Collection (ATCC), Manassas, VA, USA) prior to use as described in Malladi et al. [12], with TCID_50_ titres calculated using the method of Spearman and Kärber [14].

Identity of virus stocks were confirmed by next-generation sequencing using a MiniSeq platform (Illumina, Inc; San Diego, CA, USA). RNA was purified from Trizol-inactivated material using a Direct-zol RNA Miniprep kit (Zymo Research; Irvine, CA, USA). Purified RNA was further concentrated using an RNA Clean-and-Concentrator kit (Zymo Research). RNA was converted to double-stranded cDNA, ligated then isothermally amplified using a QIAseq FX single cell RNA library kit (Qiagen, Hilden, Germany). Fragmentation and dual-index library preparation was conducted with an Illumina DNA Prep, Tagmentation Library Preparation kit. Average library size was determined using a Bioanalyser (Agilent Technologies; San Diego, CA, USA) and quantified with a Qubit 3.0 Fluorometer (Invitrogen; Carlsbad, CA, USA). Denatured libraries were sequenced on an Illumina MiniSeq using a 300-cycle Mid-Output Reagent kit as per the manufacturer’s protocol. Paired-end Fastq reads were trimmed for quality and mapped to the published sequence for the SARS-CoV-2 reference isolate Wuhan-Hu-1 (RefSeq: NC_045512.2) using CLC Genomics Workbench version 21 from which consensus sequences were generated. Stocks were confirmed to be free from contamination by adventitious agents by analysis of reads that did not map to SARS-CoV-2 or cell-derived sequences.

### 2.5. Live virus neutralisation assays

Virus neutralisation assays (VNT) were carried out using VeroE6 cells as described previously [12]. Briefly, each serum sample was diluted 1:80 (or 1:160 where sample volume was insufficient for 1:80) in DMEM-D in a deep-well plate, followed by a two-fold serial dilution up to 1:10,240 (or 1:20,480 where 1:160 was the starting dilution). The dilution series for each serum sample was dispensed into rows of a 96-well plate, for a total volume of 50 μl per well, and triplicate wells per sample dilution. For the serum-containing wells, 50 μl virus diluted in medium to contain approximately 100 TCID_50_ (checked by back-titration) was added to each well. A positive control serum was included to confirm the reproducibility of the assay. The plates were incubated at 37°C/5% CO_2_ for 1 h to allow neutralisation complexes to form between the antibodies present in the sera and the virus. At the end of the incubation, 100 μl VeroE6 cells (2×10^4^ cells/well) were added to each well and the plates were returned to the incubator for 4 days. Each well was scored for the presence of viral CPE, readily discernible on Day 4 post-infection, with SN_50_ neutralisation titres calculated using the Spearman-Kärber formula [14] and transformed to log_2_ values for analysis. Replicates that did not show neutralisation at the lowest dilution tested (i.e., 1:80 or 1:160) were scored as ≤ 1:57 and ≤ 1:113 respectively. For statistical analysis, all samples yielding results below the detection limit were assigned a value of 1:57 based on the assumption that samples with a 1:80 starting dilution would have 100% neutralisation at one lower dilution (1:40), thereby yielding a titre of 1:57 based on the Kärber calculation (please refer to **Supplementary Text B**).

### 2.6. Modelling of SARS-CoV-2 spike protein based on in silico methods

Models of the S protein receptor binding domains (RBD, residues 332 to 532) of the major variants of concern, (Alpha, Beta, Gamma, Delta and Omicron) as well as the trimeric hCMP construct were built using AlphaFold (version 2.1.1) [15]. Models were inspected visually using VMD [16], to highlight and map the relative positions of mutations. AlphaFold provided consistent models for the RBD domains with high confidence scores, however it provided multiple conformations for the trimeric constructs.

### 2.7. Statistical analysis

VNT titre data were analysed and expressed as Log base 2 (Log_2_). Vaccine group means and standard deviations were calculated and expressed as Mean and standard deviation of mean (SD). Both one-way and two-way ANOVA was used to test the statistical differences between vaccines, antigens, and adjuvants. If the ANOVA returned a p-value of <0.05, a post hoc with Tukey’s range test (Tukey’s HSD) was performed to measure the pairwise difference and interactions between two variables. All statistical procedures were performed in R [17]) using the *car* and *lme* libraries.

## 3. Results

The Mynvax vaccine formulations elicit high antibody titres in mice that received prime-boost immunisations on Days 0,21 and 42 against the wildtype SARS-CoV-2 virus. We have previously shown that ELISA and neutralisation titres are virtually identical post the first and second boosts [11]. Neutralisation assays were performed using VIC31-D614G, Delta and Omicron SARS-CoV-2 variants. The raw data for neutralising antibody titres for individual mice as well as the mean values for the positive control serum used in the neutralisation assays are provided in **Table 1**, including the assignment of individual mice to antigen-adjuvant vaccine formulations. The box plot in **Figure 3** panels A-C represent the antibody titres for mice in each group with the median values shown as horizonal line in the plot. As with all neutralising antibody assays, a positive control serum was included in the study as assay control and back titrations were performed on the virus pools used in the assay. All control samples passed the assay criterion and virus back titrations were within the acceptable limits.

**Figure 3.**
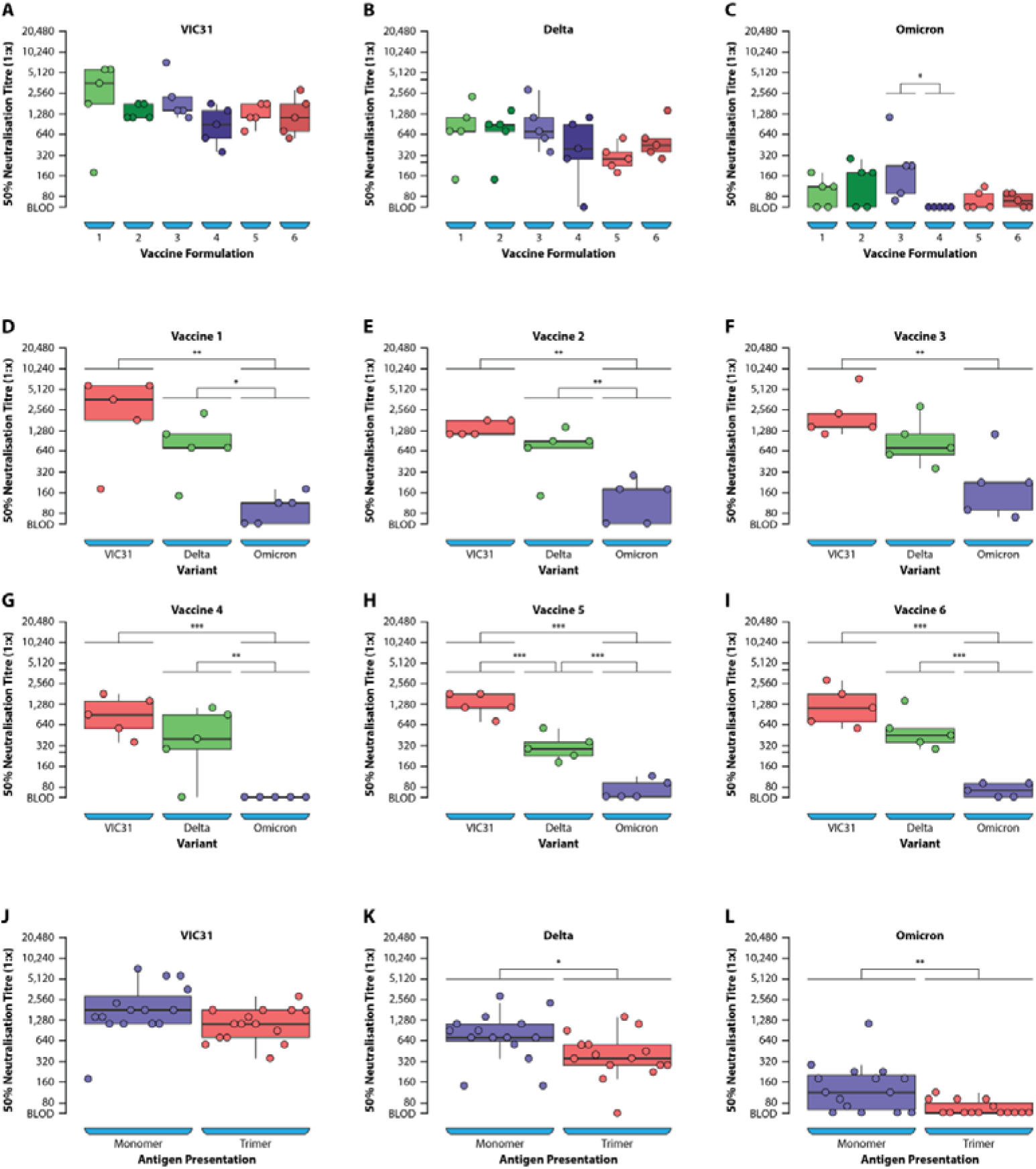
ANOVA analysis of neutralising antibody titres against SARS-CoV-2 VIC31-D614G, Delta and Omicron BA.1.1 variants following mouse immunisation with different vaccine formulations (panel A-I) and comparing monomeric to trimeric formulations (panel J-L). * p <0.05; ** p <0.001; *** p <0.001.

**Table 1.** Summary of raw data on neutralising antibody titres to VIC31-D614G, Delta and Omicron BA.1.1 SARS-CoV-2 variants for individual mice, and assignment to antigen-adjuvant vaccine formulation groups. The mean titres for the positive control serum used in the neutralisation assay are also provided. For a schematic diagram of the different vaccines 1-6 see **Figure 2**.

| Vaccine | Antigen – Presentation – Adjuvant | Mice | SARS-CoV-2 Variants* |  |  |
| --- | --- | --- | --- | --- | --- |
|  |  |  | VIC31-D614G | Delta | Omicron |
| 1 | mRBD1-3.2 – Monomer – Addavax™ | 1.1 | 1810 | 718 | 180 |
|  |  | 1.2 | 180 | 143 | ≤57 |
|  |  | 1.3 | 3620 | 718 | 113 |
|  |  | 1.4 | 5747 | 1140 | ≤57 |
|  |  | 1.5 | 5747 | 2281 | 113 |
| 2 | mRBD1-3.2 – Monomer - SWE | 2.1 | 1140 | 905 | 285 |
|  |  | 2.2 | 1140 | 143 | ≤57 |
|  |  | 2.3 | 1810 | 905 | 180 |
|  |  | 2.4 | 1810 | 718 | ≤57 |
|  |  | 2.5 | 1140 | 1437 | 180 |
| 3 | mRBD1-3.2 + mRBD1-3.2-beta – Monomer - SWE | 3.1 | 7241 | 2874 | 1140 |
|  |  | 3.2 | 2281 | 1140 | 71 |
|  |  | 3.3 | 1437 | 718 | 90 |
|  |  | 3.4 | 1437 | 570 | 226 |
|  |  | 3.5 | 1140 | 359 | 226 |
| 4 | hCMP-mRBD – Trimer – SWE | 4.1 | 570 | 285 | ≤113 |
|  |  | 4.2 | 1810 | 905 | ≤113 |
|  |  | 4.3 | 905 | 404 | ≤113 |
|  |  | 4.4 | 359 | ≤113 | ≤113 |
|  |  | 4.5 | 1437 | 1140 | ≤57 |
| 5 | hCMP-mRBD – CHO – Trimer – SWE | 5.1 | 718 | 359 | ≤57 |
|  |  | 5.2 | 1140 | 226 | ≤57 |
|  |  | 5.3 | 1140 | 180 | 90 |
|  |  | 5.4 | 1810 | 570 | 113 |
|  |  | 5.5 | 1810 | 285 | ≤57 |
| 6 | mRBD-GlyIZ – Trimer – SWE | 6.1 | 718 | 570 | 90 |
|  |  | 6.2 | 570 | 359 | 90 |
|  |  | 6.3 | 1140 | 453 | 71 |
|  |  | 6.4 | 2874 | 285 | ≤57 |
|  |  | 6.5 | 1810 | 1437 | ≤57 |
| Control Serum | Positive Control |  | 160 | 113 | 20 |

Note: \*All live virus neutralisation studies involved sera collected on Day 57 from mice vaccinated with dose 1 (day 0) dose 2 (day 21) and dose 3 (day 42) of the respective adjuvanted antigen. SN<sub>50</sub> titres are expressed as reciprocal of the neutralising titre as calculated using the Spearman-Kärber formula. Titres expressed as ≤57 and ≤113 represent analyses for which the neutralising titres were below the assay limit of detection (lowest dilution tested) of 1:80 and 1:160 respectively. For statis-
tical and fold change calculations, all $\leq 57$ and $\leq 113$ results were assigned a value of 57. See **Supplementary Text B** for further explanation on titre calculations.

The mean neutralising antibody titres (Log_2_-transformed) with standard deviations from the means, for the three variants and different vaccine formulations, are presented in **Table 2**. There was an average 2.7-fold reduction in neutralising titres to Delta compared with VIC31-D614G, whereas an average 15.4-fold reduction was noted for Omicron, across all six vaccine formulations. For monomeric formulations the fold reduction values were 2.5 and 14.4, and for trimeric 3.0 and 16.5, to Delta and Omicron respectively compared to VIC31-D614G. The fold-reduction for Omicron, compared to Delta, was 5.7- and 6.0-fold for monomeric and trimeric formulations respectively.

**Table 2.** Log_2_ transformed mean neutralising antibody titres against VIC31-D614G, Delta and Omicron BA.1.1 SARS-CoV-2 variants for mice immunised with different vaccine formulations.

| Variant | Estimate | Vaccine 1 | Vaccine 2 | Vaccine 3 | Vaccine 4 | Vaccine 5 | Vaccine 6 |
| --- | --- | --- | --- | --- | --- | --- | --- |
| VIC31-D614G | Mean | 11.02 | 10.42 | 11.02 | 9.76 | 10.29 | 10.22 |
|  | SD<sup>†</sup> | 2.09 | 0.37 | 1.07 | 0.95 | 0.56 | 0.95 |
| Delta | Mean | 9.49 | 9.36 | 9.75 | 8.52 | 8.22 | 9.02 |
|  | SD | 1.47 | 1.28 | 1.14 | 1.71 | 0.64 | 0.90 |
| Omicron | Mean | 6.56 | 6.96 | 7.69 | 5.83 | 6.16 | 6.16 |
|  | SD | 0.72 | 1.06 | 1.57 | 0.00 | 0.47 | 0.33 |

<sup>†</sup>SD = standard deviation from the mean.

The neutralising antibody titres for the three SARS-CoV-2 variants were compared for the six vaccine groups (**Figure 3A-I**). There was a reduction in neutralising titres to Delta ranging from 2.1- to 4.2-fold for the six vaccine formulations, when compared to VIC31-D614G, although this reduction was only statistically significant for Vaccine 5 (4.2-fold, p <0.01) (**Tables 2 & 3**). The reduction in neutralisation of Omicron was more pronounced, ranging from a 10.1- to 22.0-fold decrease for the six vaccine formulations, when compared to VIC31-D614G, and was statistically significant (p <0.01) for all. Vaccine 3 yielded neutralising titres 10-fold lower to Omicron when compared with VIC31, whereas the other 5 formulations yielded an average fold decrease of 16.5 (range 11- to 22-fold). The fold-reduction (range 4.2 to 7.6) for Omicron compared to Delta was also statistically significant (p <0.05) for all vaccine formulations except Vaccine 3 (p <0.1).

**Table 3.** One-way ANOVA analysis according to different variables.

| Variable | Variant of Concern | F-value | Pr(>F)<sup>a</sup> | Tukey's HSD post hoc Comparison ('adjusted' p-value) |
| --- | --- | --- | --- | --- |
| Antigen | VIC31-D614G | 1.012 | >0.100 | No significant difference |
|  | Delta | 1.463 | >0.100 | No significant difference |
| | Omicron | 3.771 | <0.050 | Antigen 2 vs Antigen 4 & 5 $p < 0.1$ |
| Adjuvant | VIC31-D614G | 1.534 | >0.100 | No significant difference |
|  | Delta | 0.688 | >0.100 | No significant difference |
|  | Omicron | 0 | >0.100 | No significant difference |
| Presentation | VIC31-D614G | 3.412 | <0.010 | Monomer vs Trimer $p < 0.1$ |
| | Delta | 4.687 | <0.050 | Monomer vs Trimer $p < 0.05$ |
| | Omicron | 5.294 | <0.010 | Monomer vs Trimer $p < 0.01$ |
| Vaccine | VIC31-D614G | 0.939 | >0.100 | No significant difference |
|  | Delta | 1.131 | >0.100 | No significant difference |
| | Omicron | 3.07 | <0.050 | Vaccine 3 vs Vaccine 4 $p < 0.05$ |
| Variant | Vaccine 1 | 10.96 | <0.010 | VIC31-D614G vs Omicron $p < 0.01$<br>Delta vs Omicron $p < 0.05$ |
| | Vaccine 2 | 16.21 | <0.001 | VIC31-D614G vs Omicron $p < 0.01$<br>Delta vs Omicron $p < 0.01$ |
| | Vaccine 3 | 8.63 | <0.010 | VIC31-D614G vs Omicron $p < 0.01$<br>Delta vs Omicron $p < 0.1$ |
| | Vaccine 4 | 13.14 | <0.010 | VIC31-D614G vs Omicron $p < 0.001$<br>Delta vs Omicron $p < 0.01$ |
| | Vaccine 5 | 68.05 | <0.0001 | VIC31-D614G vs Omicron $p < 0.0001$<br>VIC31-D614G vs Delta $p < 0.001$<br>Delta vs Omicron $p < 0.001$ |
| | Vaccine 6 | 35.66 | <0.0001 | VIC31-D614G vs Omicron $p < 0.0001$<br>VIC31-D614G vs Delta $p < 0.1$<br>Delta vs Omicron $p < 0.001$ |

a – Estimated p-value.

A summary of the One-way ANOVA analysis (**Table 3**) shows that only the “Variant” and “Presentation” variables are responsible for statistically significant differences in neutralising titres noted between different vaccine groups. No significant difference (p >0.05) was noted in neutralising antibody titres for three variants following immunisation with the different vaccine formulations, except for Omicron Vaccine 3 vs. Vaccine 4 (p <0.05) (**Figure 3A-C**; **Table 3**). The mean antibody titres to VIC31-D614G following immunisation with the vaccine formulations comprising antigens presented as monomers (Vaccines 1-3) showed a statistically non-significant (p <0.1) increase when compared to the trimers (Vaccines 4-6) (**Figure 3J-L**; **Table 3**). However, a statistically significant increase in average VNT titres was noted for both Delta (p <0.05) and Omicron (p <0.01) variants when comparing monomeric to trimeric antigen presentation (**Figure 3J-L**; **Table 3**). No significant difference (p >0.05) was noted in neutralising antibody for all three variants when comparing the six antigens or adjuvants used in vaccine formulations (**Table 3**).

In order to understand these experimental results, it was informative to look at predictions of our *in silico* molecular dynamics (**Figure 4**) and AlphaFold (**Figure 5**) modelling. As all six vaccine antigens described in this paper are RBD derivatives, the homology model in **Figure 4** shows the mutations in this domain for the five VOC to date; the difference in epitopes for each VOC is highlighted in corresponding colours. Vaccine 3 is matched to Beta, but this VOC only shares N501Y and K417N mutations with Omicron. In this study, we have used Omicron BA.1.1, which has an additional R346K mutation, along with a constellation of mutations in S (compared to past VOC, c.f. **Figure 4**).

**Figure 4.**
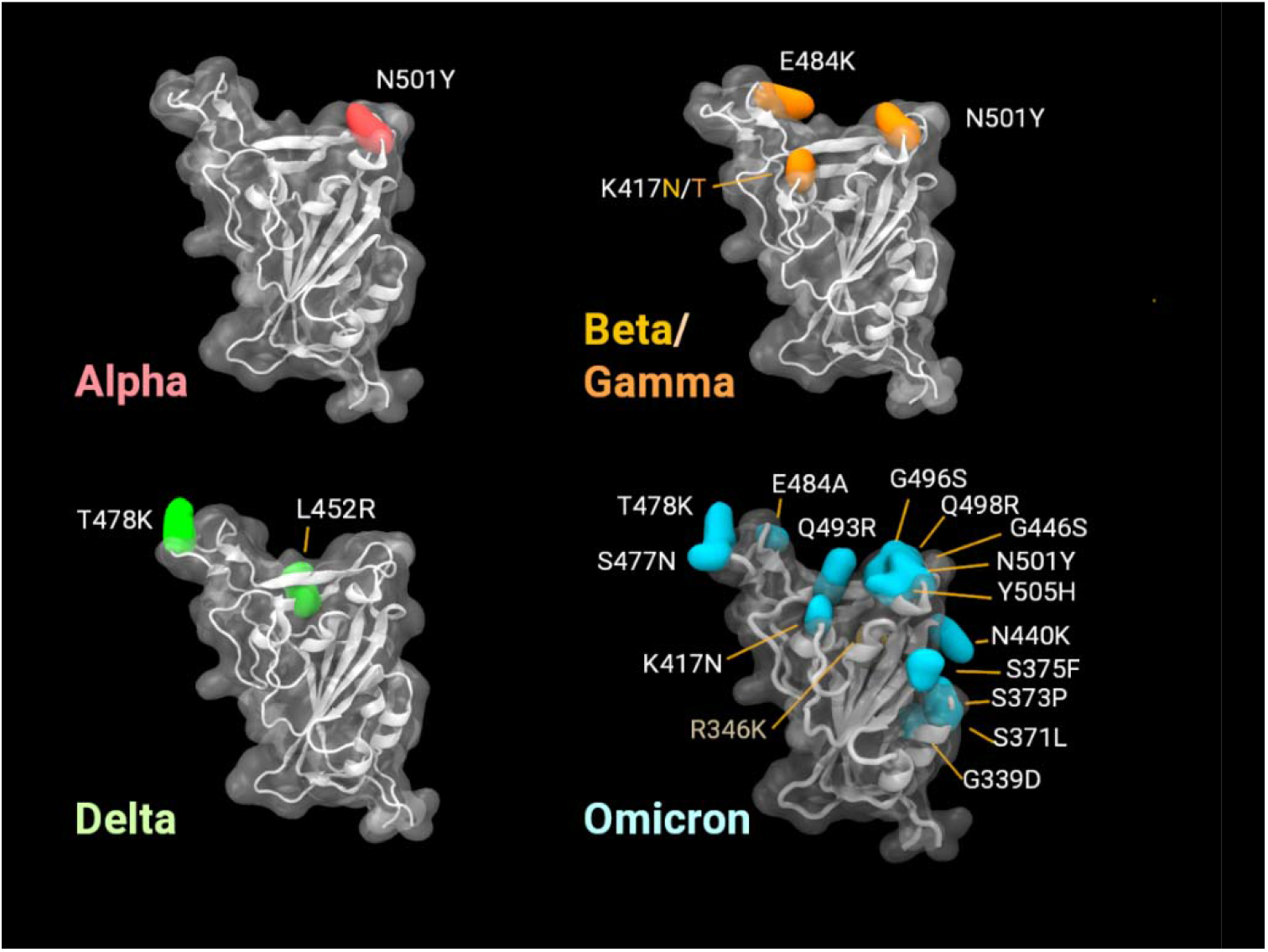
Visualisation of variant mutations in the RBD of the SARS-CoV-2 spike protein (residues 330 to 530). Omicron BA.1.1 is notable for the numbers of new mutations in this region (15 compared to 3 in Beta/Gamma and 1 in Alpha). Our isolate also included the mutation R346K. Beta and Gamma variants present similar RBDs; differing at position 417, by Asn (N).

**Figure 5.**
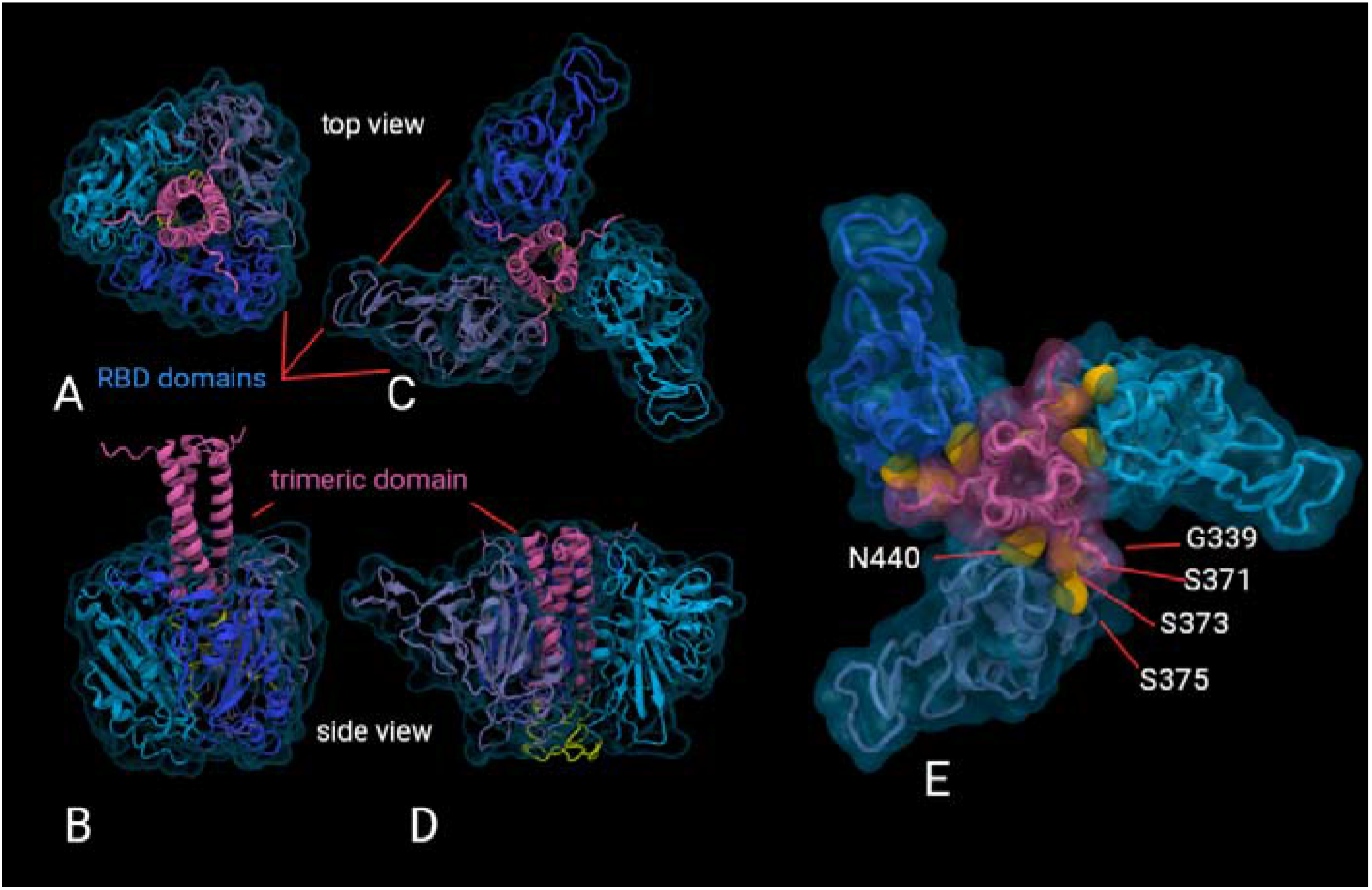
Visualisations of AlphaFold predictions of the trimeric construct. **(A)** and **(B)** show the top and side views of one probable structure; **(C)** and **(D)** of another, respectively. RBD domains are shown in blues, while the trimerisation domain is shown in pink. **(E)** shows **(C)** in greater detail, highlighting in yellow the buried G339, S371, S373, S375 and N440 residues which are mutations sites in Omicron.

AlphaFold predictions of likely structures for the trimeric formulations reveal a high probability of epitopes being buried, and inaccessible for recognition by B-cells for antibody development. **Figure 5** shows two examples of AlphaFold2 predictions of the trimeric structure, as this artificial intelligence software is currently only able to give possible structures for the multimer. That we do not have a definitive structure prediction is not a handicap for our purpose because we are able to show that key neutralising epitopes are inaccessible in each of these probable trimeric structures (showing the G339, S371, S373, S375, N440 sites mutated in Omicron VOC as an example). We also have negative stain transmission electron micrograph published elsewhere [12 (c.f. Figure 1h)] which shows that the lobed structure shown in **Figure 5C-E** is more consistent with experimental data.

## 4. Discussion

The emergence of SARS-CoV-2 VOC with considerable mutations in the spike protein, such as Omicron (BA.1.1) and waning immunity in vaccinated individuals [9,18], highlight the need for improved vaccines that are more easily deployable with fewer logistical constraints. The ability of the Omicron variant to escape neutralisation by a significant proportion of monoclonal antibody cocktails [19,20], as well as serum or plasma from vaccinated and/or infected patients [21,22], further highlights the need for a regular review of the ability of vaccine induced immunity to neutralise circulating variants, and potential subsequent vaccine matching. The development of highly thermo-tolerant monomeric and trimeric RBD derivatives [10–12] that can withstand 100 °C for 90 minutes and 37°C for four weeks have the potential to ease logistical burdens related to cold chain storage and shipping requirements of existing vaccines significantly. However, temperature stability of ‘warm’ vaccines is of little importance if they are not sufficiently Immunogenic.

In previous work [11,12], we have shown that serum samples from mice vaccinated with different formulations of this ‘warm’ vaccine were able to neutralise Alpha, Beta, Gamma, and Delta (which were the known VOC at that time). As Omicron is fast replacing all past VOC, including Delta which is still prevalent, we chose to compare neutralising antibody responses of polyclonal mouse antisera generated against these ther-motolerant vaccine candidates against Delta, Omicron, and a reference isolate used in our previous work (VIC31-D614G) [23]. In line with recent findings from studies using monoclonal antibodies, mAb cocktails or patient sera, there was a significant reduction in neutralising ability of serum collected from immunised mice against the Omicron (BA.1.1) variant, compared to VIC31-D614G and Delta. In fact, the level of neutralising antibodies to Omicron was below the detection limit of the assay in a large proportion of samples, particularly those from trimeric antigen-adjuvant formulation groups. Due to low sample volumes available from the mice in this study, the high detection limit of the VNT assay used has to be acknowledged.It appears that monomeric antigen-adjuvant formulations elicited better neutralising antibody responses compared to trimeric formulations, as demonstrated by statistically significantly higher neutralising titres to Delta and Omicron variants. These data suggest a potential advantage in pursuing the monomeric rather than trimeric formulations for Phase I clinical trials in humans. The average 14.4- or 16.5-fold reduction in neutralisation against Omicron BA.1.1 for the monomeric and trimeric formulations, respectively, compares favourably with equivalent reductions observed with leading COVID-19 vaccines [9,18]. The present neutralisation assays were carried out with sera elicited after three immunisations as sera collected after two immunisations had been exhausted. However, we have previously shown that neutralisation titres elicited after two and three immunisations with the stabilised monomeric RBDs are identical [11].

The proportional reduction in neutralising antibodies to Omicron (BA.1.1), compared to VIC31-D614G and Delta rather than complete immune evasion, suggests that mouse antibodies directed to very specific epitopes in the Spike protein of the WT virus are being evaded through mutations, while antibodies to other epitopes are still able to provide significant level of virus neutralisation. This supports previous findings showing complete evasion of specific candidate therapeutic monoclonal antibodies, and no evasion of others [19]. Therefore, a stronger polyclonal antibody response likely contributes to protection by neutralisation through sheer numbers, despite significant evasion by the Omicron variant.

In addition to efforts to induce higher antibody levels in individuals (for example through booster vaccination with homologous antigens), another approach is regular vaccine matching to current circulating variants, and/or customisation of vaccine formulations to include wild-type sequences and any relevant mutations in antigenic sites. The mRBD1-3.2 + mRBD1-3.2-beta antigen used in the formulation of Vaccine 3 in this study [10,11] was prepared using wild-type and Beta variant sequences at the time when the latter was a major concern. Although not statistically significant, there was a slightly stronger neutralisation of Omicron BA.1.1 by serum samples collected from mice immunised with this antigen, compared to only WT antigen. In fact, only in this group did all mice elicit a neutralising response above the detection limit of our assay against Omicron, whereas at least two mice from all other groups had responses below the assay detection limit. However, the high detection limit in our assay has to be noted and was due to low sample volumes available from mouse experimental serum. Nevertheless, the data suggests there might be value in adapting vaccine formulations to include key mutations from emerging variants in the spike protein of vaccine antigens. Further studies are required to investigate this and consider mutations that are common to BA.1 and BA.2, as well as those unique to each sub-lineage, given that they are antigenically distinct [24].

To understand our experimental findings, it would be beneficial to take a deeper look at the mutations in Omicron and compare monomers with trimers, as discussed below. With 15 mutations, the Omicron receptor binding domain (RBD) presents a highly altered surface compared to the Wuhan-Hu-1 reference isolate, explaining the poor neutralisation response from the vaccines based on the original strain. Even the Delta variant, with only 2 mutations in the RBD (L452R and T478K) has significantly reduced antibody neutralisation effects, suggestive of how specific vaccine-derived antibodies can be. The mutation positions of some variants are shown in Figure 4 as snapshots from our molecular dynamics simulations.

The trimeric vaccine construct is centred with a motif, namely a disulphide linked coiled-coil domain derived from the human cartilage matrix protein (residues 298-340) (hCMP) to the N-terminus of the RBD via a L14 linker [12]. The hCMP trimerisation domain is a relatively short, disulphide-linked stretch that we have previously used in the context of a safe and efficacious HIV-1 vaccine formulation in rhesus macaques [25]. We therefore also expect it to be safe in humans, though this remains to be confirmed. Of note, a Spike-derived vaccine formulation developed by Clover Biopharmaceuticals contains a much larger trimerisation domain derived from the C-propeptide of human type I(α) collagen has successfully cleared Phase 1 trials and is currently being tested in a Phase 2/3 trial [26,27]. However, the monomeric formulation of the vaccine was observed to elicit a slightly superior immune response, potentially as it presents more of antigenic epitopes, and this could be explained by the following *in silico* insights.

AlphaFold predictions of the trimeric construct rank several possible structures (as shown in **Figure 5**) with various arrangements of the RBD against the trimerised hCMP domain. One highly altered region of the Omicron BA.1.1 RBD occurs along the side face including changes G339D, S371L, S373P, S375F and N440K which occurs approximately 90 degrees orientation to the receptor binding face. This coincides with some of the predicted trimer stem/RBD interfaces of AlphaFold models. If this proves to be a stable arrangement the trimeric configuration may reduce this particular RBD epitope exposure, thus elicit a limited antibody repertoire to the full RBD. The high degree of variation in this region of the Omicron BA.1.1 RBD may contribute a significant effect to immune evasion. It remains unclear how representative the AlphaFold prediction are or how flexible these trimeric constructs (and thus how much of the epitopes are presented to the immune system) may be within a cellular environment, once administered. However, the predictions in **Figure 5E**, taken together with the transmission electron micrograph presented elsewhere [12 (c.f. Figure 1h)], leads to a likely scenario of key epitopes being buried and inaccessible for antibody development. Other possible structures, for example **Figure 5A-B**, also reveal similar stearic hindrance, therefore the lack of a definitive prediction by AlphaFold doesn’t affect our conclusions.

## 5. Conclusions

There are two opposing effects, one being that an oligomeric presentation is likely to induce higher titres of binding and neutralising antibodies, the other being steric hindrance of important epitopes required for effective B-cell responses as well as elicitation of irrelevant antibodies against the oligomerisation domain [28]. It appears that the latter effects could contribute to this host’s (mice) response to the trimeric antigen formulations; however, more studies are required with a wider range of hosts and variants before firm conclusions can be drawn. At this dosage, monomeric formulations appear relatively more effective at generating neutralising antibody responses against VOC and could be preferred for phased clinical trials of this vaccine in humans. The average 14.4-fold reduction in neutralisation against BA.1.1 observed here, compares favourably with other COVID-19 vaccines; vaccine matching should help improve the titres against targeted variants even further. The thermostability of this vaccine and its ability to withstand transient heat shocks is especially promising to address vaccine inequity that affects most low- and lower middle-income countries.

## Supplementary Materials

The following supporting information can be downloaded at: www.mdpi.com/xxx/s1, Supplementary Text A; and www.mdpi.com/xxx/s2, Supplementary Text B.

## Author Contributions

Conceptualisation, R.V. and S.S.V.; methodology, P.JvV., A.J.M., M.J.K. and N.B.S.; software, M.J.K., T.W.D. and N.B.S.; validation, S.A., M.S.K., S.K.M., R.S., S.P. and R.V.; formal analysis, N.B.S.; investigation, P.JvV., A.J.M., M.P.B., S.R., S.G., K.R.B., M.T., L.C. and J.D.D.; resources, T.W.D., R.V. and S.S.V.; data curation, N.B.S.; writing—original draft preparation, P.JvV., A.J.M., M.J.K., N.B.S., S.M., S.C., S.A. and S.S.V.; writing—review and editing, M.P.B., S.R., S.G., S.C., T.W.D., K.R.B., M.T., L.C., J.D.D., M.S.K., S.K.M., R.S., S.P. and R.V.; visualisation, A.J.M., N.B.S., M.J.K, S.M., S.C. and S.S.V.; supervision, R.V. and S.S.V.; project administration, S.C.; funding acquisition, S.S.V. All authors have read and agreed to the published version of the manuscript.

## Institutional Review Board Statement

The animal study was reviewed and approved by Institutional Animal Ethics Committee of Indian Institute of Science (IAEC no. CAF/ETHICS/799/2020).

## Data Availability Statement

The original contributions presented in the study are included in the article/Supplementary Material. Further inquiries can be directed to the corresponding author.

## Acknowledgments

We thank our colleagues from the RV Lab at IISc and its spin-out company Mynvax Private Limited for helpful discussions. The vaccine development work was supported by grants from the Bill and Melinda Gates Foundation (reference number INV-005948), and the Government of India’s Office of the Principal Scientific Advisor (reference number SP/OPSA-20-0004), and Department of Science and Technology (reference number EMR/2017/004054) to R.V., and by funding from the Australia’s Department of Finance to the CSIRO (P.I.: S.S.V.). We also acknowledge funding for infrastructural support from the following programs of the Government of India: DST-FIST, UGC Center for Advanced Study, MHRD-FAST, DBT-IISc Partnership Program, and JC Bose Fellowship to R.V. Mynvax Private Limited acknowledges funding support from IISc CSR grant for COVID-19 vaccine work, and S.S.V. acknowledges funding from National Health and Medical Research Council (grant MRF2009092) that supported growth and characterisation of variants of concern. The content is solely the responsibility of the authors and does not necessarily represent the official views of the funding or affiliated institutions.

## Supplementary Text 1

### Amino Acid Sequences of Vaccine Formulations

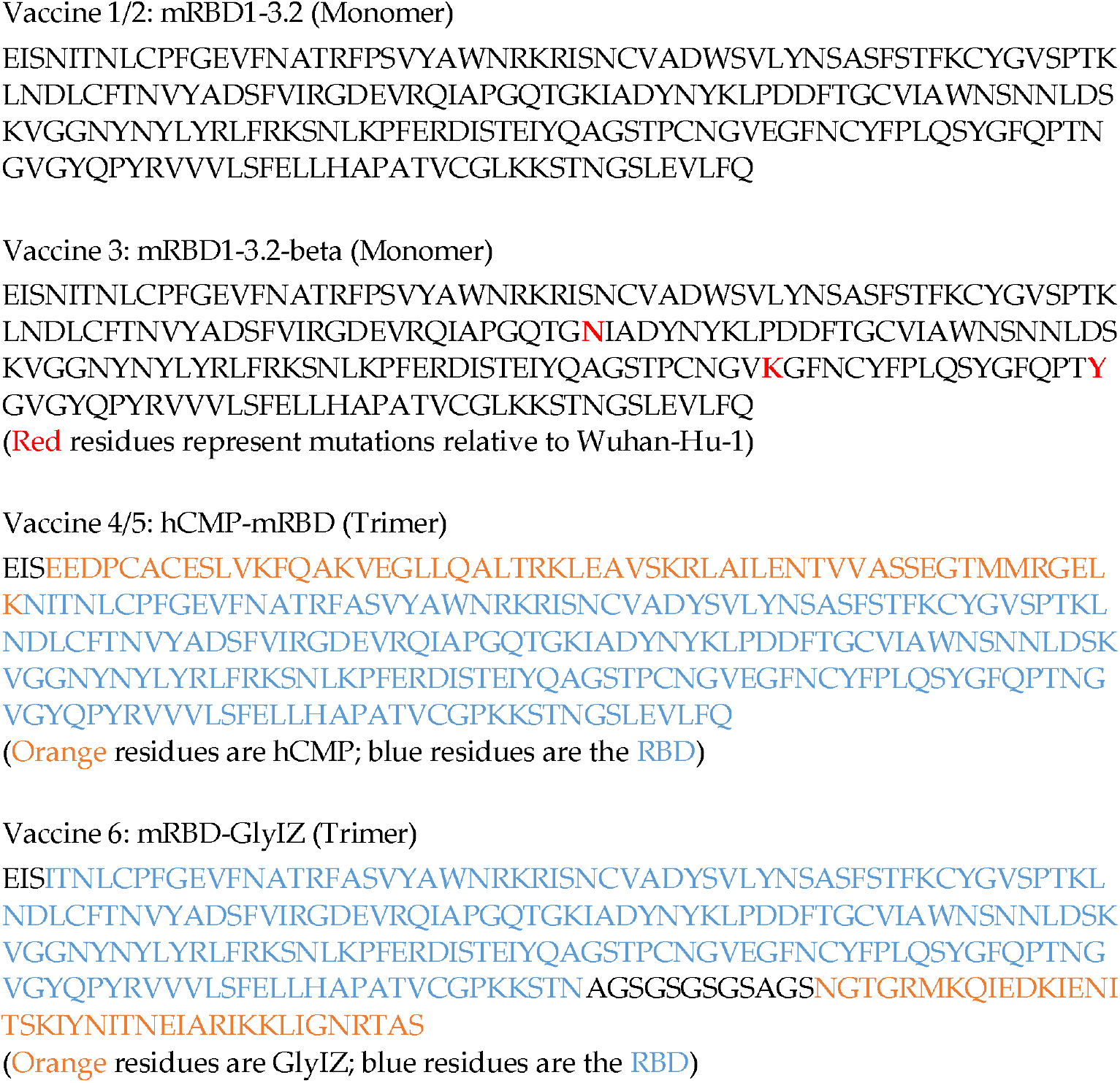

## Supplementary Text 2

### Complications in the Analysis of Microneutralisation Assay Data

#### 1. Handling of Data Below or Above the Limit of Detection (LOD)

Given that sample volumes are frequently low, especially when collected from small animal models such as mice, dilution series often need to be started at suboptimal sample dilutions. This poses problems for the analysis and interpretation of data, e.g., lack of neu-tralisation at a starting dilution of 1:80 does not mean that the sample contains no neutral-ising antibodies. Convention would have an arbitrary value assigned to the sample in this situation, however the particular arbitrary value used can substantially affect the calcula-tion of fold-differences and statistical analysis. Not having a standardised approach across different research groups will make it difficult for the WHO and other public health bod-ies to make meaningful comparisons and policy decisions [29].

The conservative approach would be to assign a value one dilution factor lower than the lower LOD (or one dilution factor higher than the upper LOD) for any sample for which a quantifiable titre cannot be obtained. Although this approach will frequently overstate neutralisation of low-responding samples, this will result in a sample below the lower LOD (BLOD) having the maximum possible neutralisation titre, and a sample above the upper LOD having the minimum possible neutralisation titre. Accordingly, calculated fold change differences between groups will be the minimum, and statistical comparison will be most conservative.

Example 1: Low sample volume availability means that four samples have to be used at a starting dilution of 1:40. Each sample is run in triplicate.

##### Results

Sample 1: 80, 160, 160
Sample 2: BLOD, BLOD, BLOD
Sample 3: BLOD, 40, 40
Sample 4: BLOD, 40, BLOD

##### Analysis

Sample 1: An NT_50_ value can be calculated using the Spearman-Karber approach as 180
Sample 2: All samples are below the LOD (no neutralisation at 1:40). Arbitrary value set at one dilution below, i.e., replicate titres 20, 20, 20. NT_50_ value calculated as 28
Sample 3: One sample below LOD, two have neutralisation at 1:40. Arbitrary value set at one dilution below, i.e., replicate titres 20, 40, 40. NT_50_ value calculated as 45
Sample 4: Two samples below LOD, one has neutralisation at 1:40. Arbitrary value set at one dilution below, i.e., replicate titres 20, 40, 20. NT_50_ value calculated as 36

#### 2. Handling of Data Below the LOD with Different Starting Dilutions

Where possible, samples should be run using the same starting dilution, however occasionally it is necessary to balance assay resolution with sample availability. If comparisons are to be made between such samples, every effort should be made to mini-mise the difference between the starting dilutions (ideally no more than one dilution fac-tor different). For titres that can be calculated from the dilution series, there will be no issues with the difference in starting dilution, however if samples have titres below the LOD there will be complications. In this situation, the arbitrary value assigned to all sam-ples below the LOD should be that of the lower starting dilution. The exception being if one or more replicates have neutralisation at the higher starting dilution. In this situation, an arbitrary value one dilution lower than that sample’s starting dilution will be used.

Example 2: Low sample volume for some of the samples means that two samples can be used at a starting dilution of 1:10, while two samples have to be started at 1:20. Each sample is run in triplicate.

##### Results

Sample 1 (starting dilution 1:10): 20, 40, 20
Sample 2 (starting dilution 1:10): BLOD, BLOD, BLOD
Sample 3 (starting dilution 1:20): BLOD, BLOD, BLOD
Sample 4 (starting dilution 1:20): 20, BLOD, 20

##### Analysis

Sample 1: An NT_50_ value can be calculated using the Spearman-Karber approach as 36
Sample 2: All samples are below the LOD (no neutralisation at 1:10). Arbitrary value set at one dilution below, i.e., replicate titres 5, 5, 5. NT_50_ value calculated as 7
Sample 3: All samples are below the LOD (no neutralisation at 1:20). Arbitrary value set at one dilution below lowest starting dilution used in assay (1:10), i.e., replicate titres 5, 5, 5. NT_50_ value calculated as 7
Sample 4: One samples below LOD, two have neutralisation at 1:20. Arbitrary value set at one dilution below sample starting dilution, i.e., replicate titres 20, 10, 20. NT_50_ value calculated as 22

